# Genome-wide association study of anti-Müllerian hormone levels in pre-menopausal women of late reproductive age and relationship with genetic determinants of reproductive lifespan

**DOI:** 10.1101/487181

**Authors:** Katherine S. Ruth, Ana Luiza G. Soares, Maria-Carolina Borges, A. Heather Eliassen, Susan E. Hankinson, Michael E. Jones, Peter Kraft, Hazel B. Nichols, Dale P. Sandler, Minouk J. Schoemaker, Jack A. Taylor, Anne Zeleniuch-Jacquotte, Deborah A. Lawlor, Anthony J. Swerdlow, Anna Murray

## Abstract

Anti-Müllerian hormone (AMH) is required for sexual differentiation in the fetus, and in adult females AMH is produced by growing ovarian follicles. Consequently, AMH levels are correlated with ovarian reserve, declining towards menopause when the oocyte pool is exhausted. A previous genome-wide association study identified three genetic variants in and around the *AMH* gene that explained 25% of variation in AMH levels in adolescent males but did not identify any genetic associations reaching genome-wide significance in adolescent females. To explore the role of genetic variation in determining AMH levels in women of late reproductive age, we carried out a genome-wide meta-analysis in 3,344 pre-menopausal women from five cohorts (median age 44–48 years at blood draw). A single genetic variant, rs16991615, previously associated with age at menopause, reached genome-wide significance at *P*=3.48×10^−10^, with a per allele difference in age-adjusted inverse normal AMH of 0.26 SD (95% CI [0.18,0.34]). We investigated whether genetic determinants of female reproductive lifespan were more generally associated with pre-menopausal AMH levels. Genetically-predicted age at menarche had no robust association but genetically-predicted age at menopause was associated with lower AMH levels by 0.18 SD (95% CI [0.14,0.21]) in age-adjusted inverse normal AMH per one-year earlier age at menopause. Our findings support the hypothesis that AMH is a valid measure of ovarian reserve in pre-menopausal women and suggest that the underlying biology of ovarian reserve results in a causal link between pre-menopausal AMH levels and menopause timing.

## Introduction

Anti-Müllerian hormone (AMH) is a member of the transforming growth factor-beta superfamily that regulates the growth and development of ovarian follicles in females and is required for sexual differentiation in the fetus, causing regression of the Müllerian ducts during testis development (1). In males, AMH is required for testes development and function and levels increase rapidly shortly after birth, peaking at 6 months of age, and then decline to low levels in during puberty (2). In women, AMH is produced by the granulosa cells of growing follicles and levels are correlated with the number of growing follicles and are used as a clinical measure of ovarian reserve (3). AMH levels increase in women from birth until their 20s, before declining gradually with age until levels are undetectable after menopause when ovarian reserve is exhausted (1, 3–5). Since AMH levels are stable throughout the menstrual cycle, they can be used as a measure of fertility in women of late reproductive age and to predict response to fertility treatment (6).

AMH levels vary widely between women and genetic variation is thought to be important, though few genetic studies have been conducted. Rare *AMH* mutations have been found with functional effects on AMH signalling (7, 8), while polymorphisms in *AMH* or the gene coding for its receptor, *AMHR2*, have been associated with response to ovarian stimulation, infertility, follicle recruitment, primary ovarian insufficiency and polycystic ovary syndrome (9).

A previous genome-wide association study (GWAS) in 1,360 adolescent males and 1,455 adolescent females from a single cohort identified three genetic variants in and around the *AMH* gene that were independently associated with higher levels of AMH in adolescent males (*P*=2×10^−49^ to *P*=3×10^−8^ for each variant when jointly included in the regression model) (10). None of these variants showed strong evidence of statistical association in adolescent females (*P*=8×10^−4^ to *P*=0.9 for each variant when jointly included in the regression model), with considerably weaker effect estimates than in males. For all three variants there was strong statistical evidence of a sex difference (*P_HET_*=3×10^−4^ to *P_HET_*=6×10^−12^), with the three cumulatively explaining 24.5% of the variation in AMH levels in males compared with 0.8% in females. No cohorts were available for replication of this initial study and it is unknown whether the weak or absent association in adolescent females persists into older ages, as would be expected since differences in ovarian decline result in variation in AMH levels between women.

We undertook a GWAS meta-analysis of 3,344 women from five cohorts – the Generations Study, Sister Study, Nurses’ Health Study, Nurses’ Health Study II and Avon Longitudinal Study of Parents and Children (ALSPAC) - to investigate genetic determinants of AMH levels in pre-menopausal women of late reproductive age (median age at blood draw 44–48 years). We aimed to identify novel genetic variants associated with AMH levels and to explore the effects of published genetic variants associated with AMH levels in previous GWAS and candidate gene studies.

## Results

### AMH is associated with a single significant signal in a known menopause locus

In our genome-wide meta-analysis (Table 1), a single genetic variant in the *MCM8* gene at 20p12.3 reached genome-wide significance at *P*<5×10^−8^ (rs16991615, *P*=3.48×10^−10^) (Figure 1, Supplementary Figure 1). Within each of the five genotyped cohorts, we inverse-normally transformed AMH and tested the association of over 11 million autosomal genetic variants imputed to HRC r1.1 2016 (11) adjusted for age and genetic relatedness (12). We performed inverse variance weighted meta-analyses of the genome-wide results from the five cohorts, filtering our results to include only variants present in three or more of the five cohorts analysed, resulting in a total of 8.4 million variants in the final results dataset. A total of 242 variants had *P*<1×10^−5^, resulting in 24 signals following distance-based clumping of variants within 500kb, with the top ten signals presented in Table 2. The minor A allele of rs16991615 increased age-adjusted inverse normal AMH by 0.26 SD per allele (95% CI [0.18,0.34], *P*=3.48×10^−10^) (Table 2).

**Table 1.** AMH levels and age of women in each of the five cohorts included in the genome-wide analysis.

| Study | n | Median (interquartile range) |  |
| --- | --- | --- | --- |
|  |  | AMH (pmol/L) | Age (years) at blood draw |
| Generations Study | 379 | 3.9 (0.8,11.7) | 44 (40,48) |
| Sister Study | 438 | 1.2 (0.1,6.0) | 48 (45,51) |
| Nurses' Health Studies (Illumina)<sup>1</sup> | 225 | 5.5 (1.5,12.4) | 45 (42,48) |
| Nurses' Health Studies (OncoArray)<sup>1</sup> | 417 | 6.7 (2.5,14.9) | 44 (41,46) |
| Avon Longitudinal Study of Parents and Children | 1,885 | 2.0 (0.4, 5.2) | 46 (44,49) |
| Total | 3,344 |  |  |

<sup>1</sup>Data from the Nurses' Health Study and Nurses' Health Study II were combined and genotyped on two different genome arrays, which were included as separate sub-samples in this analysis.

**Table 2.** Top ten signals from the genome-wide analysis of age-adjusted inverse normal AMH in pre-menopausal women.

| SNPID | Chr | Pos | EA/OA/EAF | Beta (SE) | P-value | Dir. | Het. P | Imp. qual |
| --- | --- | --- | --- | --- | --- | --- | --- | --- |
| rs16991615 | 20 | 5948227 | A/G/0.068 | 0.26 (0.04) | 3.5×10<sup>-10</sup> | +++++ | 0.14 | 0.94 |
| rs62236881 | 22 | 29450193 | A/G/0.008 | 0.85 (0.16) | 1.1×10<sup>-7</sup> | +++?? | 0.77 | 0.84 |
| rs186783371 | 5 | 88062223 | T/A/0.011 | 0.64 (0.12) | 1.4×10<sup>-7</sup> | ++++? | 0.64 | 0.61 |
| rs35829351 | 11 | 6120686 | G/A/0.449 | 0.11 (0.02) | 2.7×10<sup>-7</sup> | ++++- | 0.02 | 0.89 |
| rs10732995 | 1 | 175111574 | T/C/0.966 | 0.29 (0.06) | 3.9×10<sup>-7</sup> | +++++ | 0.86 | 0.93 |
| rs76673357 | 20 | 17831206 | G/T/0.056 | 0.22 (0.04) | 5.0×10<sup>-7</sup> | +++++ | 0.47 | 0.89 |
| rs7168070 | 15 | 93910220 | T/C/0.329 | 0.11 (0.02) | 5.5×10<sup>-7</sup> | +++++ | 0.98 | 0.89 |
| rs62237617 | 22 | 28761148 | T/C/0.011 | 0.74 (0.15) | 6.5×10<sup>-7</sup> | +++?? | 0.89 | 0.82 |
| rs141456816 | 17 | 10362307 | C/T/0.013 | 0.50 (0.10) | 6.7×10<sup>-7</sup> | ++++- | 0.44 | 0.66 |
| rs6743761 | 2 | 10393228 | A/G/0.482 | 0.09 (0.02) | 2.7×10<sup>-6</sup> | ++++- | 0.11 | 0.99 |

Beta=difference in mean age-adjusted inverse normal AMH (SD) per allele. Chr=chromosome; Dir. = positive (+) or negative (-) direction of effect in Avon Longitudinal Study of Parents and Children, Sister Study, Generations Study, Nurses' Health Studies (OncoArray), Nurses' Health Studies (Illumina) respectively, with "?" indicating that the variant was absent; EA=effect allele; EAF=weighted average effect allele frequency across the studies; Het. P=P-value from Cochran's Q-test of heterogeneity of effects across the studies; Imp. qual = mean imputation quality across the studies; Pos=position in hg19/GRCh37; OA=other allele; SE=standard error.

**Figure 1.**
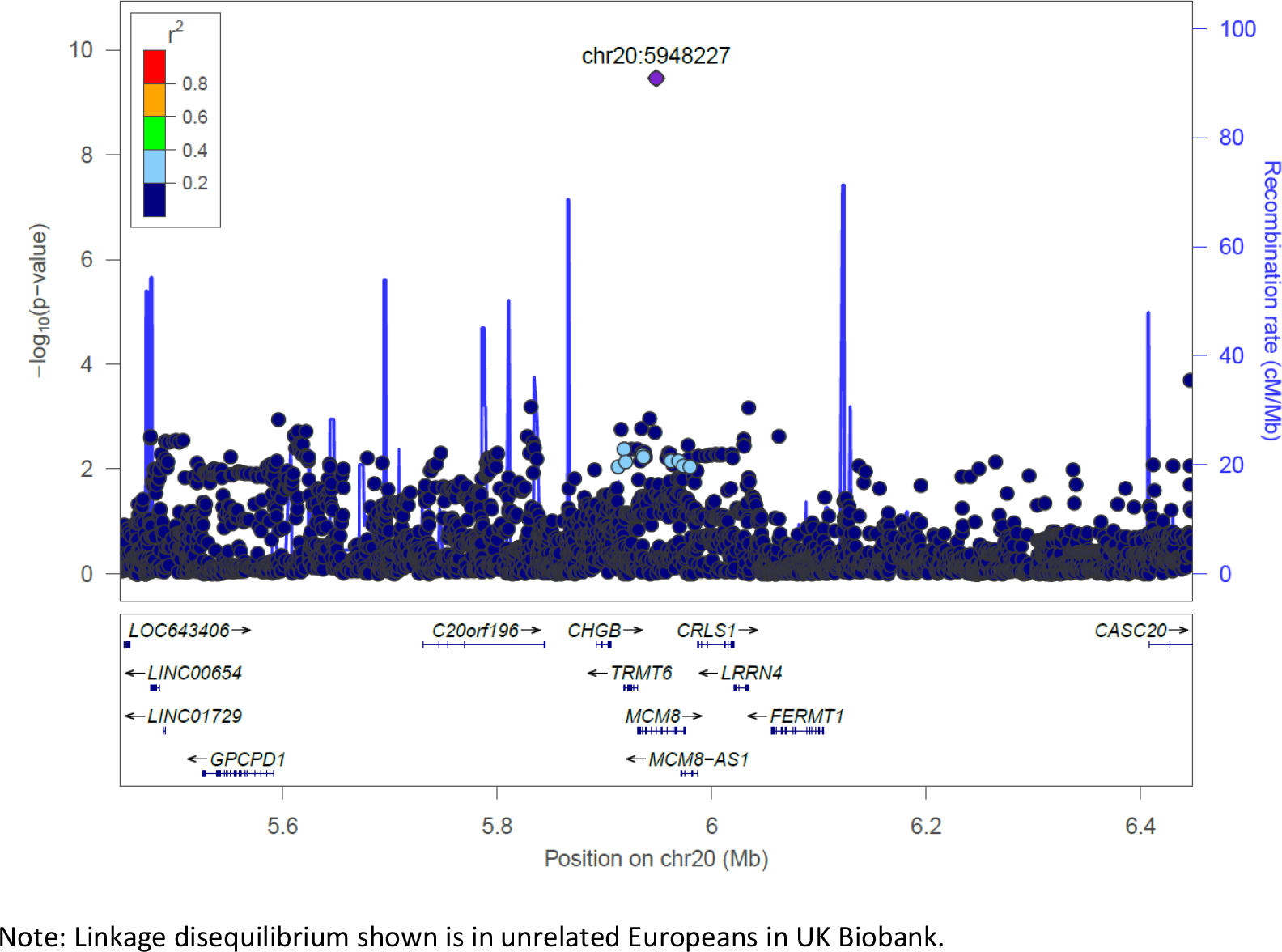
Association statistics with age-adjusted inverse normal AMH for variants within 500kb of rs16991615 (chr20:5948227) showing linkage disequilibrium with the top variant.

### Variants previously shown to be associated with AMH levels in adolescent males, but not adolescent females, were also not associated with AMH in pre-menopausal adult women in our study

For three genetic variants in and around the *AMH* gene that were previously found to be independently associated with higher levels of AMH in adolescent males (10), we estimated the effects in pre-menopausal women when the variants were jointly included in the regression model (joint model), by carrying out approximate conditional analyses using the software GCTA (13). To allow comparison between our results and the original study’s estimates, we generated effect estimates for age-adjusted inverse normal AMH in the adolescent males and females from the original study sample (ALSPAC offspring), since results from the original study were unadjusted and presented in natural log-transformed AMH.

The effect estimates from the joint model for the three published genetic variants were directionally concordant across adolescent males and females (ALSPAC offspring cohort) and the pre-menopausal women in the current study (from five cohorts including ALSPAC mothers), but had about one-fifth of the effect on the level of AMH compared with the effect in adolescent males (*PHET*<0.001) (Figure 2). The weak or null effect sizes for rs4807216, rs8112524 and rs2385821 were similar in adolescent and pre-menopausal females (*PHET*>0.05 for all). Genetic variant rs2385821 had the strongest effects in females of the three variants from the previous publication (10), but did not reach genome-wide significance in pre-menopausal females (for joint model, per allele difference in age-adjusted inverse normal AMH of 0.27 SD (95% CI [0.13,0.41]), *P*=4.0×10^−5^) (Supplementary Table 1).

**Figure 2.**
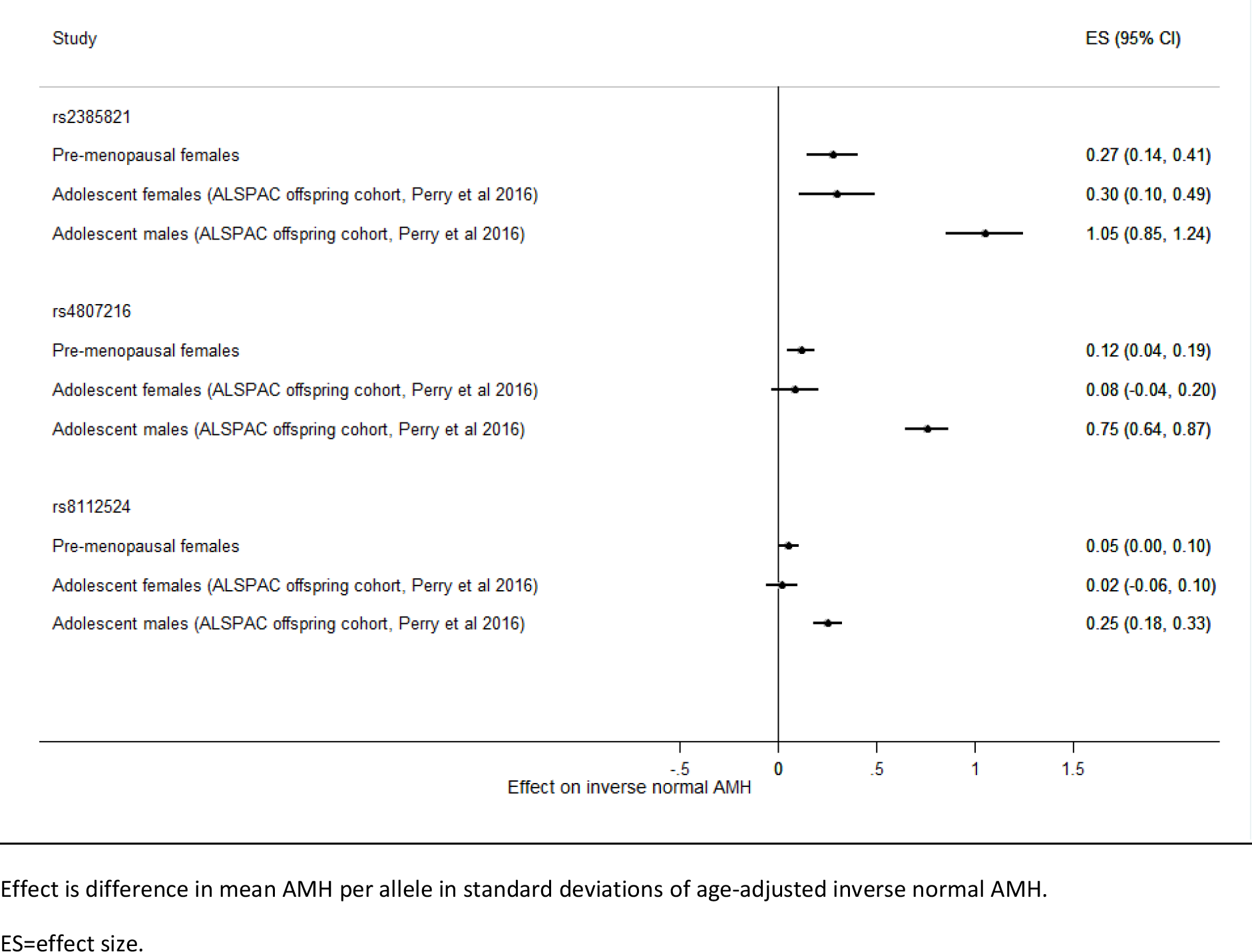
Comparison of effect sizes of three genetic variants previously associated with higher levels of AMH (10) when jointly included in the regression model: effect in adolescent males and females from the ALSPAC offspring cohort (previous study, Perry et al 2016 (10)) and pre-menopausal women (current study).

### Polymorphisms in AMH linked to reproductive phenotypes were associated with differences in AMH levels

Two non-synonymous polymorphisms in *AMH* that have been previously reported as associated with follicular phase oestradiol (14) and primary ovarian insufficiency (7) had weak effects on AMH levels. Variant rs10407022 (I49S), associated with raised follicular phase oestradiol levels (14), was associated with raised AMH levels in our study (per allele difference in age-adjusted inverse normal AMH of 0.08 SD (95% CI [0.02,0.14]), *P*=2.8×10^−3^) (Supplementary Table 2). A variant associated with premature ovarian insufficiency, rs10417628 (A515V), was associated with lower AMH levels (per allele difference in age-adjusted inverse normal AMH of −0.20 SD (95% CI [−0.35,−0.04]), *P*=0.01) (7). In contrast, rs2002555, an *AMHR2* polymorphism (−482A>G) associated with raised oestradiol levels (14) and three variants in *ACVR1* shown to affect AMH levels in women with polycystic ovary syndrome (PCOS) (rs1220134, rs10497189 and rs2033962) (15), were not robustly associated with AMH levels in our study (*P*>0.05) (Supplementary Table 2).

### Genetic variants for early menopause are associated with reduced levels of AMH

Since the only genetic variant to reach genome-wide significance in our study (rs16991615) has previously been reported as associated with menopause timing (16–18), we investigated the association of AMH levels with all 56 genetic variants associated with menopause timing (18). The effect estimates of these genetic variants on AMH level were positively correlated with the published effects on age at menopause (r=0.83) and there were consistent directions of effect for 50/56 variants (*P*=1×10^−9^ for binomial sign test; *χ*^2^_56_ =194.39, *P*=4×10^−17^ for global chi-squared test of association) (Figure 3, Supplementary Table 3). There were no obvious outliers among the 56 menopause timing variants and generally variants with large effects on age at menopause also had large effects on AMH levels.

**Figure 3.**
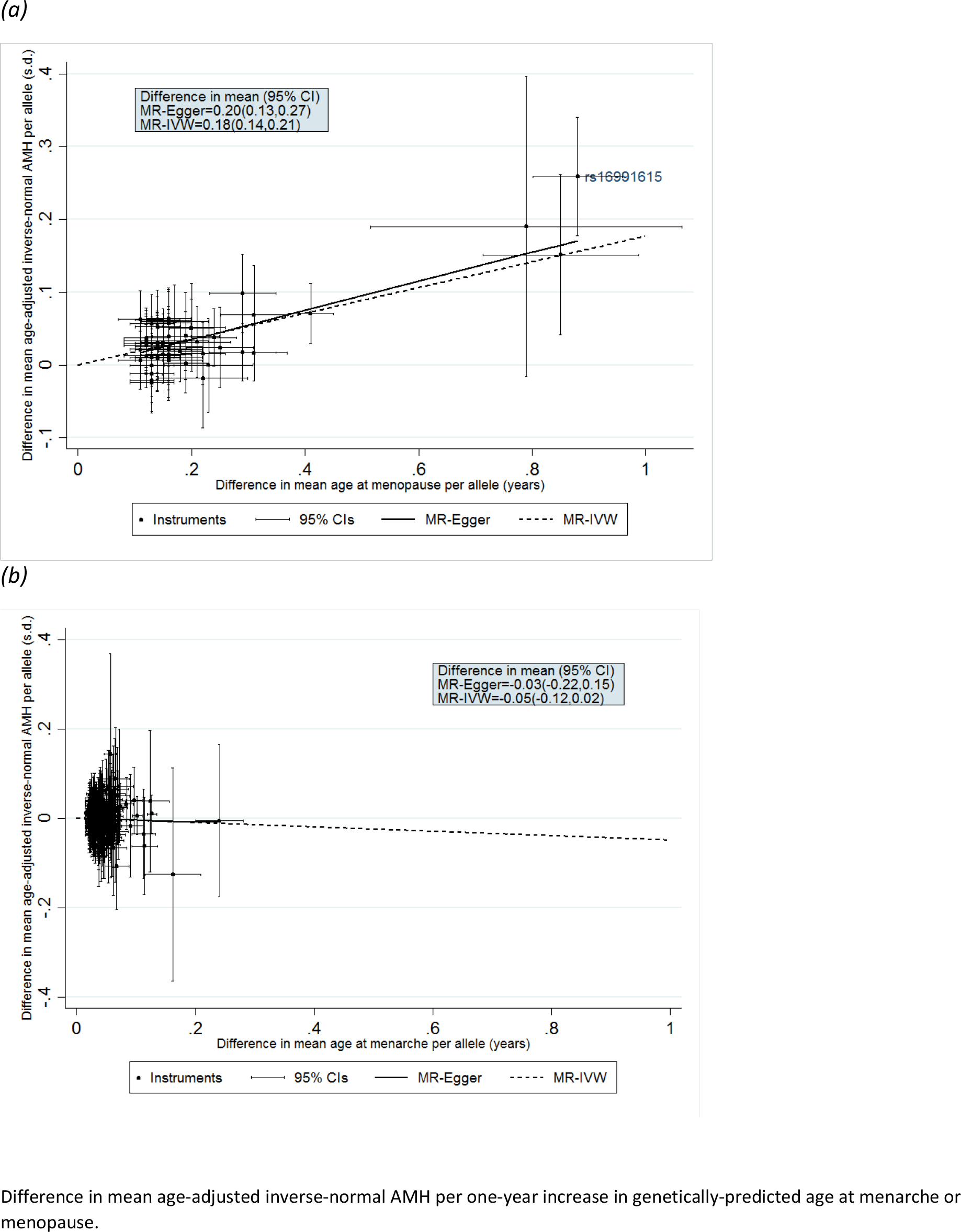
Inverse variance weighted and Egger two-sample Mendelian randomization analyses of the effect of genetically-predicted (a) age at menopause and (b) age at menarche on age-adjusted inverse normal AMH levels in pre-menopausal women.

Two-sample Mendelian randomization analysis by inverse variance weighted (IVW) and Egger estimation supported a causal relationship between genetically-predicted age at menopause and pre-menopausal AMH level (Figure 3, Supplementary Table 3). For a one-year increase in genetically-predicted age at menopause, age-adjusted inverse normal AMH was increased by 0.18 SD (95% CI [0.14,0.21]) with no horizontal pleiotropy detected by Egger analysis (in Egger analysis, 0.20 SD (95% CI [0.13,0.27]) age-adjusted inverse normal AMH per a one-year increase in genetically-predicted age at menopause one-year increase in genetically-predicted age at menopause, *P-intercept*=0.49). This relationship remained similar even when rs16991615 was excluded from the analysis (Supplementary Table 5).

### Genetic variants for age at menarche are not associated with AMH levels

We investigated the effect of genetic variants associated with age at menarche on AMH levels (19), since menarche marks the age at which ovarian reserve starts to decrease through ovulation. We identified 327 of 389 published independent signals (19) in our meta-analysis results. For the 327 variants, there was little correlation between the published effect on age at menarche and the effect on AMH levels (r=−0.05) and the directions of effects were not consistent, with 158/327 (48%) in the same direction (*P*=0.58 for binomial sign test; χ^2^_327_=328.62, *P*=0.46 for global chi-squared test of association) (Figure 3, Supplementary Table 4). Two-sample Mendelian randomization analysis by IVW and Egger estimation found no causal relationship between age at menarche and AMH level (difference in mean age-adjusted inverse normal AMH per one-year increase in genetically-predicted age at menopause was for IVW, −0.05 SD (95% CI [−0.12,0.02]) and for Egger, −0.03 SD (95% CI [− 0.22,0.15], *P-intercept*=0.87) (Figure 3, Supplementary Table 5).

### Genetic variant for follicle-stimulating hormone levels is not associated with AMH levels

Since levels of follicle-stimulating hormone (FSH) and luteinising hormone (LH) rise around menopause, we tested the association of a genetic variant at the *FSHB* locus that affects levels of these hormones (20) with AMH levels in pre-menopausal women. The *FSHB* promoter polymorphism (rs10835638; −211G>T) was not associated with AMH levels (per allele difference in age-adjusted inverse normal AMH of 0.01 SD (95% CI [−0.05,0.07]), *P*=0.79).

### Sensitivity analysis

Results for genetic variants with *P*<5×10^−5^ in the main analysis were well-correlated whether we adjusted for age or not (*r*=0.99) (Supplementary Figure 2), used our favoured inverse normal transformation or a natural log transformation (as in reference (10)) (*r*=1.00) (Supplementary Figure 3), and when we excluded women whose AMH level was imputed as it was below the lower limit of detection (*r*=0.99) (Supplementary Figure 4).

## Discussion

Our results indicate that variation in AMH levels in pre-menopausal women is contributed to by the underlying biology of ovarian reserve, as shown by the correlation between genetically-predicted age at menopause and AMH levels, supporting the use of AMH as a means of measuring ovarian reserve. The only signal passing genome-wide significance in our analyses was rs16991615 in *MCM8*, a published menopause timing variant (16–18), with the same allele associated with earlier menopause and lower AMH levels. Genome-wide analyses of menopause timing, a proxy measure for ovarian reserve, have identified 56 genetic variants and highlighted the importance of DNA damage response pathways during follicle formation *in utero* and for follicle maintenance during a woman’s lifetime (18). Additionally, 389 genetic variants have been identified for menarche timing, the age at which ovarian reserve starts to decrease through ovulation (19). Therefore, it is plausible that genetic determinants of menarche and menopause timing could affect ovarian reserve and influence AMH levels in pre-menopausal women, many years prior to menopause. For genetic variants associated with age at menopause, the published effect estimates were positively correlated with the effects on AMH levels and there was evidence of a causal relationship between genetically-predicted earlier menopause and lower pre-menopausal AMH levels, which remained even when rs16991615 was excluded. We did not find any association between genetically-predicted age at menarche and pre-menopausal AMH levels. We interpret these results as suggesting that AMH levels in pre-menopausal women are determined by declining ovarian reserve as a result of reproductive ageing but not menarche timing, and that women with lower AMH are nearer to the end of their reproductive lifespan.

Variant rs16991615 has previously been found to be associated with differences in menopause timing by 0.9 years per allele and is a missense variant in exon 9 of *MCM8* (E341K), which is required for homologous recombination (21). Other mutations in *MCM8* causing reduced double strand break repair have been found in women with premature ovarian failure (22) and follicle development is arrested at an early stage in *MCM8* knockout mice (23). Pathway analysis has shown that the menopause timing variants identified from genome-wide analyses are enriched for genes involved in DNA damage response, including double-strand break repair during meiosis, suggesting that the genetic determinants of age at menopause act during ovarian follicle formation or maintenance, potentially affecting ovarian reserve from before birth until menopause (19). Therefore, it seems likely that rs16991615 affects AMH levels through differences in ovarian reserve.

Although the three published GWAS signals in and around the *AMH* gene (10) did not reach genome-wide significance when jointly included in the regression analysis, they did show directional consistency and were nominally associated in the pre-menopausal women in our study. The previous GWAS of adolescent females included fewer samples (n=1,455) than our analysis (n=3,344), hence we were better powered to detect the effects of these variants in females. The three published variants for AMH had smaller effects in pre-menopausal women compared with adolescent males but were consistent with the associations seen previously in adolescent females. The strongest signal in the GWAS of AMH levels in adolescents, rs4807216, was not associated with age at menopause in the most recent genome-wide meta-analysis (per allele difference in age-adjusted inverse normal AMH of 0.05 SD (95% CI [-0.05, 0.15]), *P*=0.37) (18). Differences in genetic regulation of AMH levels in males and females are plausible given AMH’s different function in men and women. In males, AMH is required for regression of the Müllerian ducts during testis development in the fetus, and is involved in testicular development and function (2). In females, AMH is produced by granulosa cells of primary, pre-antral and small antral follicles, inhibiting both the further recruitment of primordial follicles from the follicle pool and also FSH-dependent selection of follicles for growth during the menstrual cycle (3, 6, 14, 24). AMH expression starts *in utero* at 36 week’s gestation, peaking at around 25 years before declining to undetectable levels at menopause (3, 5, 6). In our analysis, a polymorphism in the promoter of *FSHB* (−211G>T) that affects FSH levels (25–27) had no effect on AMH levels in pre-menopausal women, supporting the absence of direct negative feedback of FSH on AMH.

Our study provides some evidence for the effect on AMH levels of two previously identified polymorphisms in *AMH* associated with oestradiol levels (14) and premature ovarian insufficiency (7). We found no evidence for a more general role in regulation of AMH levels for variants in *ACVR1*, encoding ALK2 involved in AMH signalling, which are associated with AMH levels in women with PCOS (15). We were unable to test other genetic variants in *AMH* with functional effects on AMH signalling as they either could not be identified as a genetic variant in Build 37 of the Human Genome Assembly (28) or were not present in our final GWAS results (7).

AMH levels vary widely between women, reflecting factors such as variation in ovarian reserve, age and ethnicity (12). We controlled for age and ethnicity by adjusting for age and restricting our analyses to genetically European individuals. Adjustment for age will remove a source of variation in AMH level, the effect of the decrease in the primordial follicle pool with age, highlighting the effect of genetic variants responsible for variation in the initial size of the primordial follicle pool or that either accelerate or protect against loss of ovarian reserve with age.

We would have been unable to detect low frequency variants or those with a smaller effect size since we were only powered (>80%) to detect a variant with a MAF of 5% and an effect of 0.36 SD or greater in the sample size analysed. We were unable to evaluate the effect of time from study participation to menopause on our results, to investigate whether the association of the menopause variants was modified by proximity to menopause, since we did not have sufficient follow-up data. Future analysis should consider stratifying by participants’ time to menopause. However, such analyses would require large numbers of women who had pre-menopausal AMH measurements (AMH levels are generally undetectable post-menopause), recorded age at menopause and varying times of follow-up since age at menopause. We are not aware of any such study currently but with continued follow-up of the women included in this study such analyses may be possible.

This study confirms genetically that AMH levels are a marker for ovarian decline and reproductive ageing in pre-menopausal women. In addition to its use as a marker of fertility, there is evidence that AMH is a biomarker of breast cancer risk in pre-menopausal women. In a recent study, odds of pre- and postmenopausal breast cancer were 60% higher in women in the highest quartile of AMH level compared with the lowest, even after adjusting for potential confounders such as age (29). Our study suggests that these effects could be mediated through preserved ovarian reserve, or a correlate of ovarian reserve, as a result of delayed reproductive ageing, supported by findings from a large scale genomic analysis that showed a causal effect of later menopause on increased risk of breast cancer by 6% per year (18) and strong epidemiological evidence that later age at menopause increases risk of breast cancer (30). Our study provides genetic evidence that underlying biological factors responsible for reproductive ageing contribute to AMH levels in pre-menopausal women and are likely to be the main driver for the observed associations of AMH with health outcomes.

## Materials and Methods

### Studies included

The central analysis team at University of Exeter Medical School coordinated data collection from the five studies. We included 3,344 women who had pre-menopausal AMH levels measured, who were participants in the Generations Study (31), the Sister Study (32), the Nurses’ Health Study, the Nurses’ Health Study II (33) and ALSPAC (34–36) (Table 1) (Supplementary Information). For the Generations Study, the Sister Study and the Nurses’ Health Studies, genotype and phenotype data were provided to the central analysis team for quality control, cleaning and analysis. For the ALSPAC study, quality control and genotype-phenotype analyses were undertaken in house and summary descriptive, GWAS and sensitivity analyses statistics were provided to the central analysis team for meta-analysis.

### Genetic data

In the Generations Study, Sisters and the Nurses’ Health Studies, samples were genotyped on the OncoArray array (Table 1). For the Nurses’ Health Studies, a further 225 samples were genotyped on an Illumina array. For each cohort and array type, data were cleaned using a standard quality control process in PLINK v1.9 (www.cog-genomics.org/plink/1.9/) (37). SNPs were removed if they were poorly genotyped (missing in >5% samples) or were not in Hardy-Weinberg equilibrium (P<1×10^−6^). Samples were removed if they were poorly genotyped (missing >5% SNPs), were a sex mismatch or were outliers in terms of heterozygosity. Within each cohort, samples that were related to each other as 3rd degree relatives or closer were identified and the sample with the greater proportion of missing SNPs was removed. Principal component analysis was carried out in FlashPCA in order to identify and remove genetically non-European samples from the analysis. SNPs with MAF>1% aligned to the correct strand in HRC v1.1 were used for imputation. Genotypes for chromosomes 1– 22 were phased using SHAPEIT and imputed to HRC r1.1 2016 using the University of Michigan Imputation Server (https://imputationserver.sph.umich.edu/) (11, 38, 39).

ALSPAC mothers were genotyped using the Illumina Human660W-Quad array at Centre National de Génotypage (CNG) and genotypes were called with Illumina GenomeStudio. Quality control was performed in PLINK v1.07 (40) by removing poorly genotyped SNPs (missing in >5% samples), not in Hardy-Weinberg equilibrium (P<1×10^−6^), or that had MAF<1%. Samples were removed if they were poorly genotyped (missing >5% SNPs), had indeterminate X chromosome heterozygosity or extreme autosomal heterozygosity. Samples showing evidence of population stratification were identified by multidimensional scaling of genome-wide identity by state (IBS) pairwise distances using the four HapMap populations as a reference, and then excluded (IBS>0.125). Haplotypes were estimated using SHAPEIT (v2.r644) (41) and phased haplotypes were then imputed to HRC panel (11) using IMPUTE V3.

### AMH phenotype

For the Generations Study, Sister Study and the Nurses’ Health Studies, AMH levels were measured in blood samples taken from pre-menopausal women before breast cancer incidence by the individual studies as part of a collaborative, prospective study of AMH and breast cancer risk (29). Serum AMH levels were measured using an ultrasensitive ELISA (Ansh Labs, Webster, TX) (Sister Study) or a picoAMH enzyme-linked immunoabsorbent assay (Ansh Labs, Webster, TX) (Generations Study and the Nurses’ Health Studies, and samples below limit of detection of ELISA in Sister Study) (29). In ALSPAC, blood samples were taken following a standardized protocol in women who attended a series of clinic assessments starting about 18 years after the index pregnancy and fasted (overnight or a minimum of 6 hours for those assessed in the afternoon) serum AMH levels were measured using the Beckman Coulter AMH Gen II ELISA assay (35, 36).

For samples with AMH below the lower limit of detection, levels were imputed: for the Generations Study, the value was the midpoint between zero and the lower limit of detection (0.00821 pmol/L); for the Sister Study, missing values were imputed as 0.0015 ng/mL to be consistent with the previous analysis (29); for ALSPAC, measured AMH values <0.01 ng/mL were imputed to be 0.01 ng/mL. A small number of women (n=24) in the Nurses’ Health Studies with AMH below the lower limit of detection (2.038 pg/mL) were excluded from the analyses. For all studies, measured values of AMH were converted to pmol/L using 1 pg/mL=0.00714 pmol/L and 1 ng/mL= 7.14 pmol/L. AMH was transformed by inverse normal transformation, in which the rank of the AMH value was converted to the z-score for the corresponding quantile of a standard normal distribution, in order to approximate a normal distribution to ensure normality of the residuals in the association analysis. AMH levels for each study are summarised in Table 1.

### Genome-wide analysis

Genome-wide linear regression analysis was carried separately for each of the five cohorts for autosomal genetic variants with imputation quality>0.4 assuming an additive model. Age at time the blood sample was taken was included as a covariate since age is known to be strongly associated with AMH levels and was negatively correlated with AMH level in exploratory analysis (median age in each study is summarised in Table 1). For the Generations Study, Sisters and Nurses’ Health Study, analysis was carried out using GEMMA 0.94.1 (42), which calculates a genetic relationship matrix to account for cryptic relatedness and population stratification between the samples. The genetic relationship matrix was created from a pruned list of uncorrelated SNPs created in PLINK 1.9 www.cog-enomics.org/plink/1.9/) (37) using --indep-pairwise, excluding regions of long range linkage disequilibrium, based on variants with MAF>0.01, excluding variants with r^2^>0.5 (window size of 1000, calculated in steps of 50). The analysis included approximately 11.8 million genetic variants for the Generations Study, 12.9 million for the Sister Study, 12.3 million for Nurses’ Health Study OncoArray and 11.1 million for Nurses’ Health Study Illumina. For ALSPAC, the analysis was carried out in SNPTESTv2.5 (43) adjusting for the top ten principal components of ancestry which resulted in approximately 14.7 million SNPs.

Standard error weighted meta-analysis of the individual GWAS results was carried out in METAL (version 2011-03-25) (44) with genomic control applied to account for inflation due to any remaining population stratification. Genetic variants included in the meta-analysis had imputation>0.4 and minor allele count>5 (calculated from allele frequencies), resulting in a total of 11.2 million autosomal SNPs that were analysed. Approximately 8.4 million variants were present in three or more of the five datasets analysed and were included in our final results. We identified independent signals as being suggestive of genome-wide association if they had *P*<1×10^−5^ and were more than 500kb from another signal; from these, we identified signals reaching genome-wide significance at *P*<5×10^−8^.

Manhattan and quantile-quantile plots for the genome-wide association results were created using the package *qqman* (45) in R (The R Foundation for Statistical Computing). LocusZoom v1.4 (46) was used to plot the association statistics with age-adjusted inverse normal AMH for variants within 500kb of the top variant, showing linkage disequilibrium. Linkage disequilibrium was calculated in PLINK v1.9 (www.cog-genomics.org/plink/1.9/) (37) from best guess genotypes for 1000 Genomes Phase 3/HRC imputed variants in ̃340,000 unrelated Europeans from the UK Biobank study (47).

### Generation of age-adjusted inverse normal effect estimates in ALSPAC offspring cohort

For three published genetic variants that were associated with AMH levels in adolescent males in the ALSPAC offspring cohort (10), we generated effect estimates for age-adjusted inverse normal transformed AMH in the original study sample, since the original published estimates were unadjusted and presented in natural log-transformed AMH. Analyses were carried in SNPTEST v2.5 (43) adjusting for the top ten principal components of ancestry and age and excluding the most extreme 1% of measured AMH values, resulting in 1,312 males and 1,421 females from the ALSPAC offspring cohort for analysis. Other methods were as described previously (10).

### Estimation of joint effects of variants in and around the AMH gene

We used GCTA (version 1.25.0), using the command--cojo-joint (13), to carry out an approximate conditional analysis to estimate the joint effects of three genetic variants in and around the *AMH* gene (10). Linkage disequilibrium between the variants was estimated using a random sample of 8,569 white British individuals from the UK Biobank May 2015 interim release of imputed genetic data (48).

### Comparison of effects for published traits with AMH results

We compared the published effect estimates for 56 genetic variants associated with age at menopause (18) and 389 genetic variants associated with age at menarche (19) with their effects on AMH level in our analysis, by carrying out a Binomial sign test of directional consistency and calculating Pearson correlation coefficients. To explore whether age at menopause or age at menarche causes differences in AMH levels, we used the genetic variants associated with menopause and menarche timing as instruments for age at menopause and age at menarche in two-sample Mendelian Randomization analyses. We used the Stata package *mrrobust* (49) to carry out inverse variance weighted (IVW) and Egger (which takes account of horizontal pleiotropy (50)) analyses. Analyses were carried out in Stata MP 13.0 and Stata SE 14.2 (StataCorp, TX, USA).

### Sensitivity analysis

The genome-wide analysis was repeated without adjustment for age, using a natural log transformation (to be consistent with the previously published GWAS (10)), and also excluding women whose AMH level was imputed as it was below the lower limit of detection. We compared effect sizes in the main analysis with estimates from these alternate analyses for genetic variants with *P*<5×10^−5^ in the main analysis.

## Supporting information

## Acknowledgements

We thank Breast Cancer Now and the Institute of Cancer Research for support and funding of the Generations Study, and the study participants, study staff, and the doctors, nurses and other health care providers and health information sources who have contributed to the study. The Institute of Cancer Research acknowledges National Health Service funding to the NIHR Biomedical Research Centre.

We would like to thank the participants and staff of the Nurses’ Health Study and the Nurses’ Health Study II.

This research has been conducted using the UK Biobank Resource. We would like to thank Dr R.N. Beaumont and Dr S.E. Jones (Genetics of Complex Traits, University of Exeter Medical School) for assistance with calculating linkage disequilibrium in UK Biobank.

We are extremely grateful to all the families who took part in the ALSPAC study, the midwives for their help in recruiting them, and the whole ALSPAC team, which includes interviewers, computer and laboratory technicians, clerical workers, research scientists, volunteers, managers, receptionists and nurses.

