## Supplementary material for "Genome-wide association study of anti-Müllerian hormone levels in pre-menopausal women of late reproductive age and relationship with genetic determinants of reproductive lifespan"

**Supplementary Information**

*Details of studies included*

ALSPAC

The ALSPAC study (34–36) is a prospective population-based birth cohort study that recruited 14,541 pregnant women resident in the South West of England with expected dates of delivery from 1st April 1991 to 31st December 1992 (http://www.alspac.bris.ac.uk.). The women and their offspring have been followed-up since that date and information presented here is from a subgroup of the original mothers who were pre-menopausal at the time of AMH blood sampling (34–36). Ethical approval for the study was obtained from the ALSPAC Ethics and Law Committee and the Local Research Ethics Committees. Please note that the study website contains details of all the data that is available through a fully searchable data dictionary and variable search tool: http://www.bristol.ac.uk/alspac/researchers/our-data.

Generations Study

The Generations Study (31) is a prospective population cohort study started in 2003 to investigate the environmental, behavioural, hormonal and genetic causes of breast cancer (31). The cohort includes over 110 000 women aged 16 and older at entry, recruited from the general UK population through connections to the charity Breakthrough Breast Cancer (now Breast Cancer Now) or who volunteered as a result of publicity, and female friends and family members of participants. Follow-up questionnaires are mailed to participants about every 3 years. The study received appropriate ethical approval from the South East MREC, and informed consent was received from the participants. Detailed menstrual histories were collected and blood samples were contributed by 92% of participants.

Nurses' Health Study and Nurses' Health Study II

In 1976, 121,701 female, registered nurses, ages 30 to 55 years, were enrolled in the Nurses' Health Study (33). Biennially, participants complete mailed questionnaires on lifestyle, diet, reproductive history, and disease diagnoses. In 1989–1990, 32,826 women ages 43 to 69 years (21% premenopausal) donated blood samples.

The Nurses' Health Study II was established in 1989, when 116,430 female registered nurses, ages 25 to 42 years, completed and returned a questionnaire (33). The cohort has been followed biennially following the methods of the NHS. Between 1996 and 1999, 23,393 premenopausal participants, who were cancer-free and between the ages of 32 and 54 years, provided blood samples.

Sister Study

The Sister Study prospective cohort was designed to address genetic and environmental risk factors for breast cancer. During 2003-2009, 50,884 U.S. and Puerto Rican women ages 35-74 were recruited through a national multi-media campaign and network of recruitment volunteers, breast cancer professionals and advocates. Eligible women had a sister who had been diagnosed with breast cancer but did not have breast cancer themselves. This research was approved by the Institutional Review Boards of the National Institute of Environmental Health Sciences, NIH, and the

Copernicus Group. All participants provided informed consent. Data analysed in this study were from a subgroup of participants with a serum sample who were premenopausal (32).

### Supplementary Tables and Figures

*Supplementary Table 1. Comparison of effect sizes from univariate analyses and joint analyses (approximate conditional analyses in GCTA) in pre-menopausal women and adolescent males and females for three genetic variants associated with higher levels of AMH in adolescent males* (10).

| | | | | | Univariate analysis | | GCTA joint model | |
|---|---|---|---|---|---|---|---|---|
| | SNPID | Chr | Pos | EA/OA/EAF | Effect (SE) | *P* | Effect (SE) | *P* |
| Adolescent males | rs4807216 | 19 | 2248683 | C/T/0.135 | 0.64 (0.04) | 4.0E-47 | 0.75 (0.06) | 2.3E-39 |
| | rs2385821 | 19 | 2120154 | G/A/0.965 | 0.18 (0.09) | 0.04 | 1.05 (0.10) | 3.9E-26 |
| | rs8112524 | 19 | 2250528 | G/A/0.373 | 0.40 (0.03) | 1.3E-35 | 0.25 (0.04) | 1.9E-11 |
| Adolescent females | rs4807216 | 19 | 2248683 | C/T/0.13 | 0.01 (0.05) | 0.85 | 0.08 (0.06) | 0.18 |
| | rs2385821 | 19 | 2120154 | G/A/0.963 | 0.21 (0.09) | 0.01 | 0.30 (0.10) | 2.7E-03 |
| | rs8112524 | 19 | 2250528 | G/A/0.371 | 0.02 (0.04) | 0.66 | 0.02 (0.04) | 0.65 |
| Pre-menopausal females | rs4807216 | 19 | 2248683 | C/T/0.138 | 0.08 (0.03) | 5.2E-03 | 0.12 (0.04) | 1.5E-03 |
| | rs2385821 | 19 | 2120154 | G/A/0.963 | 0.13 (0.06) | 0.02 | 0.27 (0.07) | 4.0E-05 |
| | rs8112524 | 19 | 2250528 | G/A/0.373 | 0.07 (0.02) | 3.1E-03 | 0.05 (0.03) | 0.03 |

Effect is in difference in mean AMH per allele in standard deviations of age-adjusted inverse normal AMH.
Chr=chromosome; EA=effect allele; EAF=mean effect allele frequency; OA=other allele; SE=standard error.

*Supplementary Table 2. Association of* AMH, AMHR2 and ACVR1 *polymorphisms reported by candidate gene studies with age-adjusted inverse normal AMH in pre-menopausal women.*

| Polymorphism | Study | Chr:Pos (b37) | SNPID | EA | OA | EAF | Beta | SE | *P* |
|---|---|---|---|---|---|---|---|---|---|
| *AMHR2* -482A>G | Kevenaar ME et al 2007 (14) | 12:53817237 | rs2002555 | G | A | 0.18 | 0.01 | 0.03 | 0.67 |
| *AMH* I49S T>G | Kevenaar ME et al 2007 (14) | 19:2249477 | rs10407022 | G | T | 0.17 | 0.08 | 0.03 | 2.8E-03 |
| *AMH* A515V C>T | Alvaro Mercadal B et al 2015(7) | 19:2251817 | rs10417628 | T | C | 0.02 | -0.20 | 0.08 | 0.01 |
| *ACVR1* rs1220134 T>A | Kevenaar ME at al 2009 | 2:158603556 | rs1220134 | A | T | 0.27 | 0.01 | 0.02 | 0.81 |
| *ACVR1* rs10497189 T>C | Kevenaar ME at al 2009 | 2:158624804 | rs10497189 | C | T | 0.1 | 0.03 | 0.03 | 0.39 |
| *ACVR1* rs2033962 G>T | Kevenaar ME at al 2009 | 2:158638441 | rs2033962 | A | C | 0.18 | 0.00 | 0.03 | 0.86 |

Beta is in difference in mean AMH per allele in SD of age-adjusted inverse normal AMH.
Chr=chromosome; EA=effect allele; EAF=mean effect allele frequency; Pos=position in hg19/GRCh37; OA=other allele; SE=standard error.

*Supplementary Table 5. Results of Mendelian Randomization analyses of the effect of genetically-predicted age at menopause and age at menarche on age-adjusted inverse normal AMH levels in pre-menopausal women.*

| Exposure | Analysis | Effect (95% CI) | *P* | *P-intercept* |
|---|---|---|---|---|
| Age at menopause | IVW | 0.18 (0.14,0.21) | 9.9E-26 | n/a |
| | Egger | 0.20 (0.13,0.27) | 9.1E-08 | 0.49 |
| Age at menopause, excluding rs16991615 | IVW | 0.16 (0.12,0.20) | 1.5E-21 | n/a |
| | Egger | 0.13 (0.05,0.22) | 2.2E-03 | 0.49 |
| Age at menarche | IVW | -0.05 (-0.12,0.02) | 0.17 | n/a |
| | Egger | -0.03 (-0.22,0.15) | 0.72 | 0.87 |

Effect is difference in mean AMH in standard deviations of age-adjusted inverse normal AMH per one-year increase in age at menopause/menarche.
IVW=inverse variance weighted estimation; SE=standard error.

*Supplementary Figure 1. (a) Manhattan and (b) QQ plot for GWAS of age-adjusted inverse normal AMH in pre-menopausal women.*

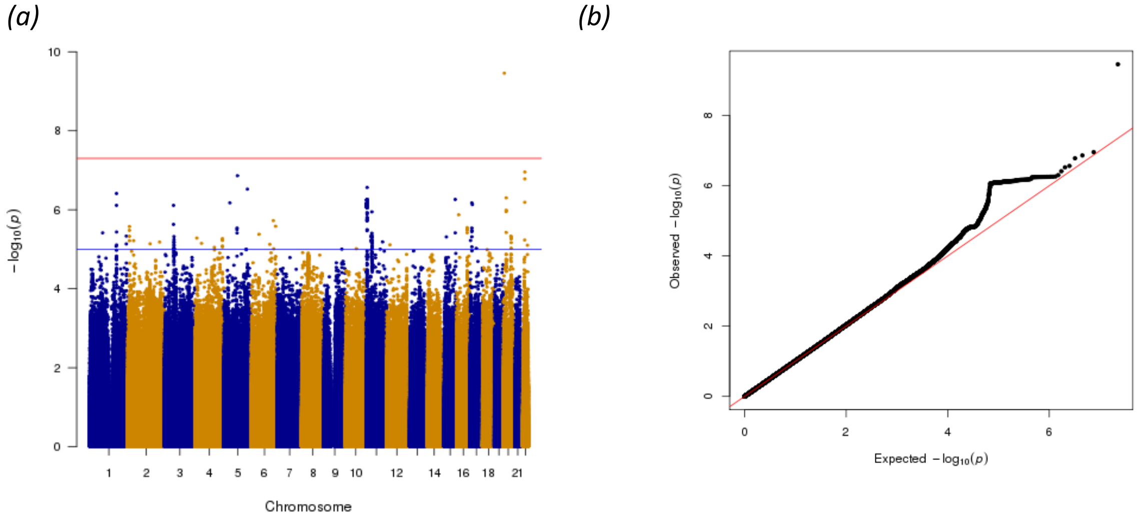


*Supplementary Figure 2. Comparison of effect sizes in the main GWAS (SD of age-adjusted inverse normal AMH) and the analysis not adjusted for age (SD of inverse normal AMH) for genetic variants that were* P<5×10$^{-5}$ *in the main GWAS.*

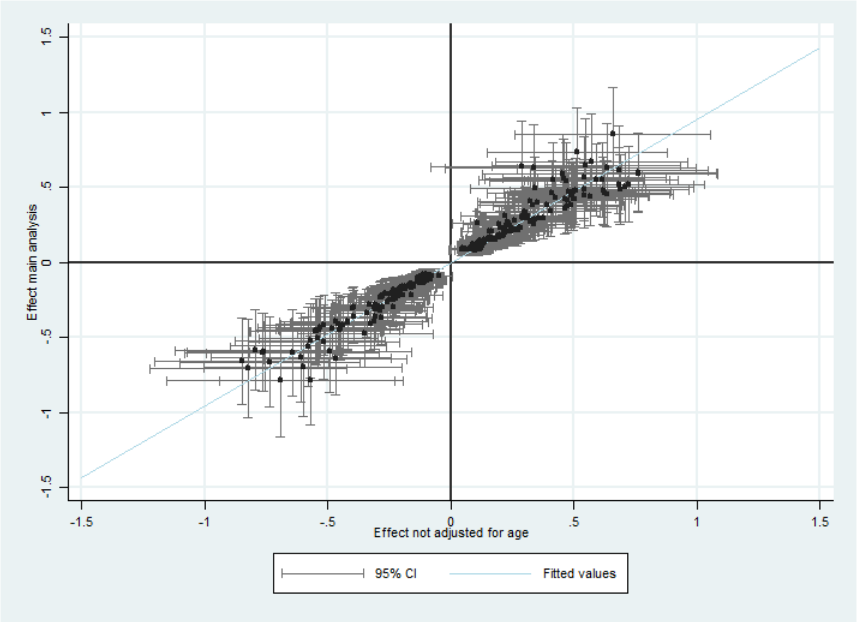

*Supplementary Figure 3. Comparison of effect sizes in the main GWAS (SD of age-adjusted inverse normal AMH) and the natural log transformed analysis (SD of age-adjusted natural log transformed AMH) for genetic variants that were* P*<5×10$^{-5}$ in the main GWAS.*

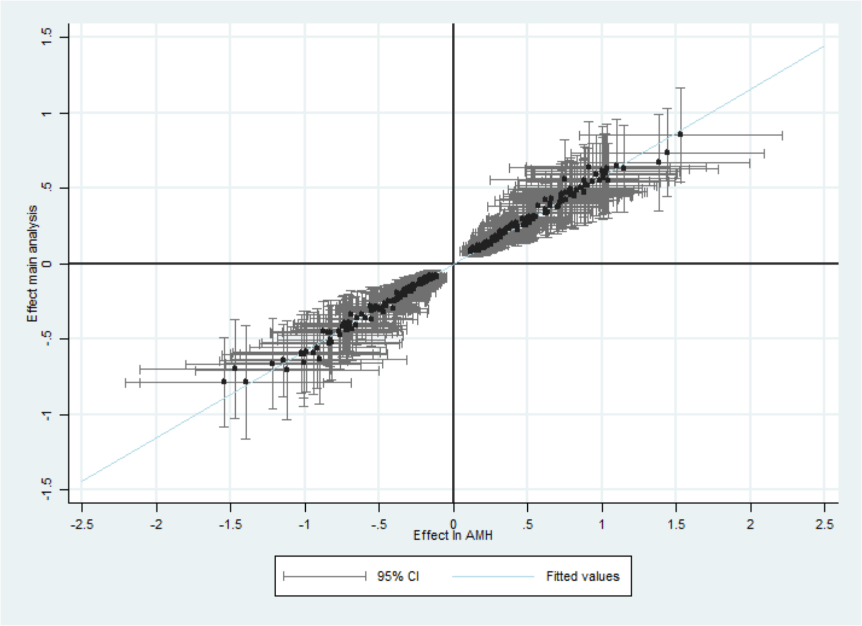


*Supplementary Figure 4. Comparison of effect sizes in the main GWAS and the analysis excluding women with AMH measured as below the lower limit of detection (effects in SD of age-adjusted inverse normal AMH for both) for genetic variants that were* P*<5×10$^{-5}$ in the main GWAS.*

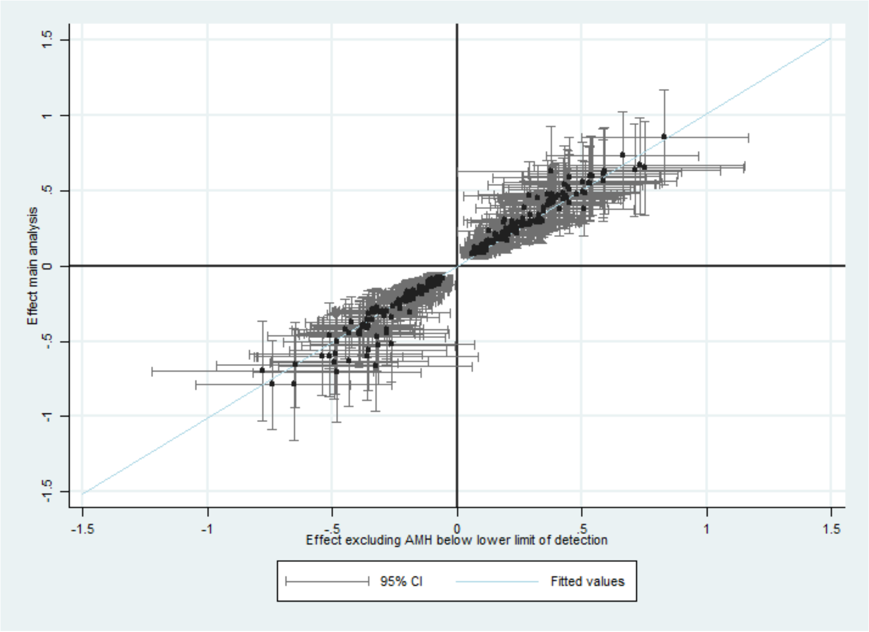
